# Dirty mice better recapitulate key features of mRNA vaccine immunogenicity observed in humans

**DOI:** 10.64898/2026.03.07.709392

**Authors:** Beatriz Praena, Frances K. Shepherd, Cera A. McDonald, Devyani Joshi, Sneh Lata Gupta, Madison L. Ellis, Lilin Lai, Alberto Moreno, Shanley N Roach, Mark J. Pierson, Adam J. Gilbertsen, Carl Davis, Mehul S. Suthar, Jens Wrammert, Ryan A. Langlois

## Abstract

Although specific pathogen free (SPF) mice have traditionally been used to test candidate vaccines, recent work has demonstrated that “dirty” mice with broad microbial exposure more appropriately recapitulate human immune responses. Using a model where lab mice are co-housed with pet store mice, we modeled SARS-CoV-2 mRNA vaccine responses in dirty and traditional SPF models. We found that dirty mice show reduced serum spike-binding antibody titers after prime and require a second booster dose to reach SPF-level spike antibody titers. Additionally, spike antibodies showed faster waning in dirty mice through 5 months post-vaccination, and neutralizing activity of these antibodies were reduced against Omicron variants, directly comparable with observations in humans. We further investigated the seasonality and consistency of pathogens in cohoused mice, and the impact of serial microbial exposure on our animal model system. We found that pathogen exposure and T cell activation remained consistent over time, and that a single co-housing event was sufficient to provide broad microbial exposure. This work demonstrates that the dirty mouse co-housing system is a promising, translationally representative approach to screen candidate mRNA vaccines for efficacy and durability prior to human clinical trials.

**Significance statement:** The development of mRNA vaccines during the COVID-19 pandemic dramatically reduced hospitalization and death rates for infected individuals. However, booster vaccinations were required to achieve full efficacy, and protection waned over time. Our research leveraged a “dirty” mouse model to test whether SARS-CoV-2 mRNA vaccinations in animals with previous microbial exposure better modelled human immune responses. We found that dirty mice require a booster vaccination for full efficacy and experienced waning serum antibody titer over time. We propose this approach as a future model for robust preclinical mRNA vaccine testing.

## Introduction

Preclinical vaccination studies are typically performed on specific pathogen free (SPF) animals raised in barrier facilities to reduce confounding variables and increase replicability. However, a consensus has begun to form that “dirty” animals with broad microbial exposure better recapitulate adult human immune activity and vaccine response (1-7). Multiple systems (3) have been designed to provide microbial exposure to laboratory mice, including co-housing with pet store mice to generate dirty mice (1), “rewilding” mice in outdoor-mimicking environments (8), “feralizing” mice through reintroduction to an outdoor environment with wild-caught mice (9), or laboratory mice born to wild mice (“wildlings”) which maintain the natural microflora (10), among others. Vaccination studies performed in SPF and dirty mice reliably show that microbial exposure affects humoral immunity (2, 5, 7). This corresponds with human data demonstrating that vaccination response in individuals from different countries and with different microbial exposures resulted in altered vaccine outcomes (11, 12). As humans naturally experience diverse microbial interactions, it follows that vaccination research intended for human treatment should consider pre-existing immune responses.

In this study, we investigated the humoral response of SPF and dirty co-housed mice to SARS-CoV-2 mRNA vaccination. While mRNA vaccines are effective, waning immunity and viral evolution require boosters (13-15). We hypothesized that dirty mice would better recapitulate this human experience than SPF mice, which overestimate immunogenicity. Indeed, dirty mice required a booster to achieve SPF-level anti-Spike IgG titers, which then waned more rapidly. Furthermore, serum from vaccinated SPF animals better neutralized SARS-CoV-2 USA-WA1/2020 and detected Omicron variants (BA1.1/BA1.5) as compared to dirty mice. Comparing mice co-housed once versus sequentially, we found a single co-housing event sufficient to recapitulate the human response. Finally, our model demonstrated consistent pathogen exposure and T cell activation over years. These results suggest dirty mice better reflect basal human immune status and therefore are a translationally representative preclinical model for evaluating vaccine modalities.

## Results

To investigate the humoral immune response to mRNA vaccines in the context of previous infection history, we vaccinated SPF and dirty mice with either Pfizer-BioNTech SARS-CoV-2 mRNA vaccine or mRNA encoding the receptor-binding domain (RBD) derived from the WA1 variant (Fig. 1A). This experiment was performed four times to ensure that observed results were consistent between the unique microbial exposures achieved through co-housing. A booster vaccination was performed 31 days (d31) after the prime vaccination. To determine if immune experience impacts antibody production after mRNA SARS-CoV-2 vaccination, serum IgG was evaluated for SARS-CoV-2 spike binding by ELISA (enzyme-linked immunosorbent assay). We observed a greater IgG response to the prime in SPF mice, which increased further after the booster. In contrast, dirty mice required a booster to reach a peak IgG response comparable to the SPF animals (Fig. 1B). Antibody titer declined between 62- and 151-days post-vaccination at a slightly higher rate in the dirty compared to SPF mice, though the difference was not significant (Fig. 1C). Further, serum antibodies from vaccinated SPF mice better neutralized SARS-CoV-2 USA-WA1/2020 30 days after booster, as determined by a reduction neutralization test (Fig. 1D), and recognized Omicron variants BA.1 and BA.5. (Fig. 1E). We additionally injected mouse IgG antibody targeting the influenza A nucleoprotein (α’NP) into the tail veins of SPF and dirty mice and observed faster reduction of α’NP antibody from the serum of dirty mice than SPF mice (Fig. 1F), suggesting that a higher antibody clearance rate in dirty mice may act as a potential mechanism for reduced vaccine titers.

**Fig. 1.**
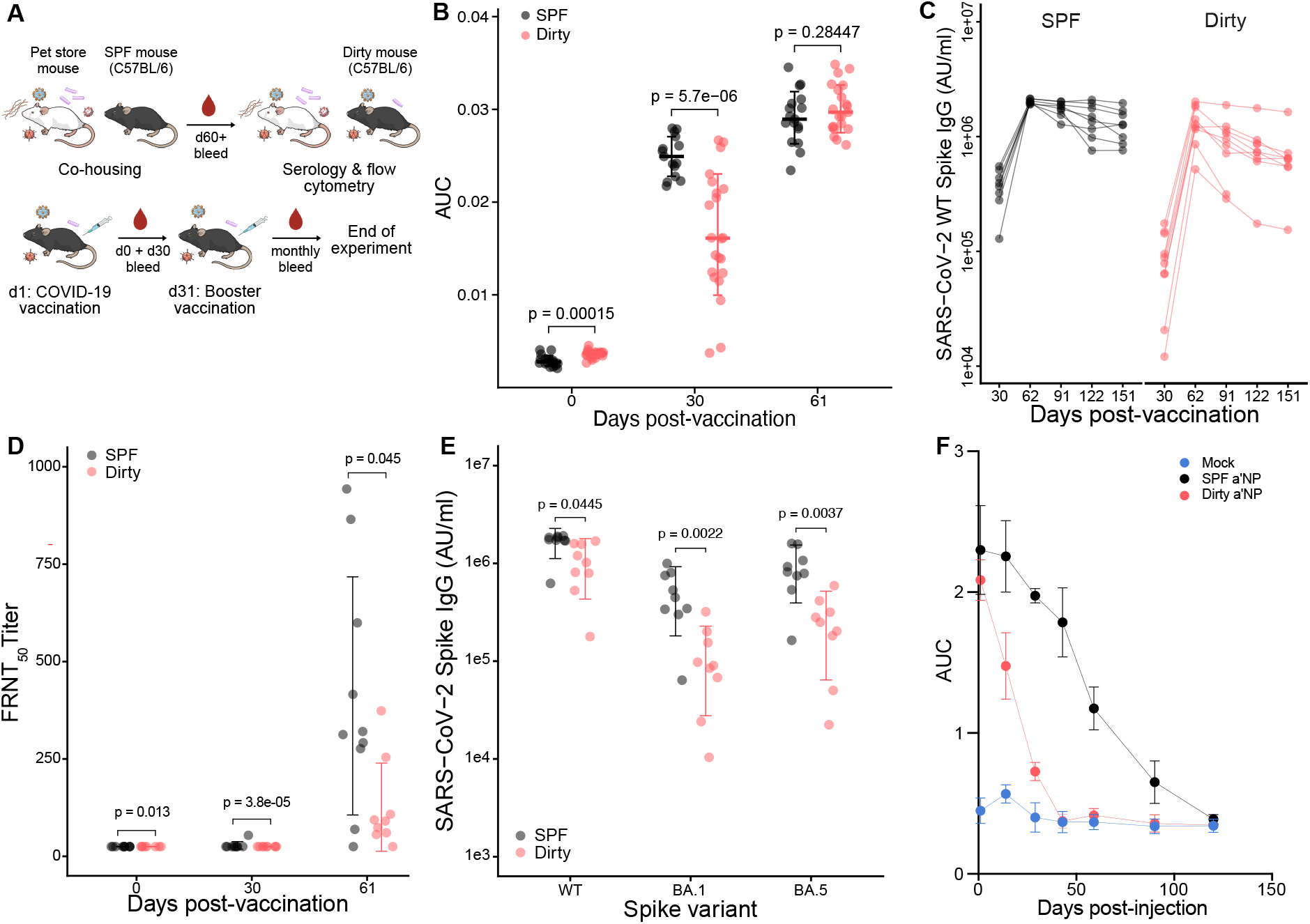
mRNA vaccination in dirty mice requires booster vaccination and is less durable than in SPF mice. (A) Model for generating and vaccinating dirty mice, days (d) indicated. (B–E) Serum antibody responses in SPF (black) and dirty (red) mice. Illustration generated with images from NIAID NIH BioArt. (B) Anti-S1 RBD IgG levels over 61 days post-vaccination (N=48), expressed as area under the curve (AUC) from serial dilution ELISAs (OD_405_). (C) Quantification of serum IgG specific to SARS-CoV-2 WT Spike (AU/ml) through day 151 post-immunization. (D) Neutralizing antibody titers over 61 days post-vaccination, expressed as FRNT_50_. (E) Cross-reactive serum IgG levels against SARS-CoV-2 Spike variants (WT, BA.1, BA.5) at day 61 (AU/ml). Data represent individual mice with mean ± SEM; p-values determined by Welch’s T-test^2^. (F) Antibody clearance from serum of mock-injected (N=3) and α’NP-injected SPF (N=4) and dirty mice (N=4), expressed as α’NP IgG2a AUC. N=3-4 AUC values per timepoint are represented on the graph aside from mock d59 with N=2, with standard deviation indicated. Antibody levels in B, C, and E were determined using MSD-ECLIA. Antibody levels in F were determined with ELISA.

We next assessed the pathogen burden in these animals and found diverse pathogens across four independent experiments (Fig. 2A). We measured the frequency of CD8a+/CD44+ T cells in the blood as a readout of immune system activation (2) and found that higher levels of activated T cells correlated with attenuated vaccine responses (p<0.001, Fig. 2B). We further evaluated the pathogen heterogeneity of the co-housing system across eight years and all four seasons using data from 1014 laboratory mice co-housed with pet store mice. We used multidimensional scaling to compare similarity of pathogen exposure profiles of the mice, revealing heterogeneity in exposure (Fig. 2C). However, overall T cell activation status was not different over time or season (Fig. 2D), suggesting the dirty mouse model is a highly robust model for immune system activation despite potential differing pathogen exposures. To investigate the effect of repeated microbial exposures on immune response, we established a double co-housing model where SPF mice were co-housed with a pet store mouse for 60 days and then re-housed with a new pet store mouse for an additional 60 days (Fig. 2E). Despite re-exposure to many pathogens (Fig. 2F), double co-housing did not significantly increase the T cell activation compared to single co-housing (Fig. 2G), and there was no difference in post-booster antibody responses to the mRNA vaccine between single and double co-housing (Fig. 2H). However, the primary response was notably more heterogeneous in double co-housed mice, consistent with their more complex microbial exposure.

**Fig. 2.**
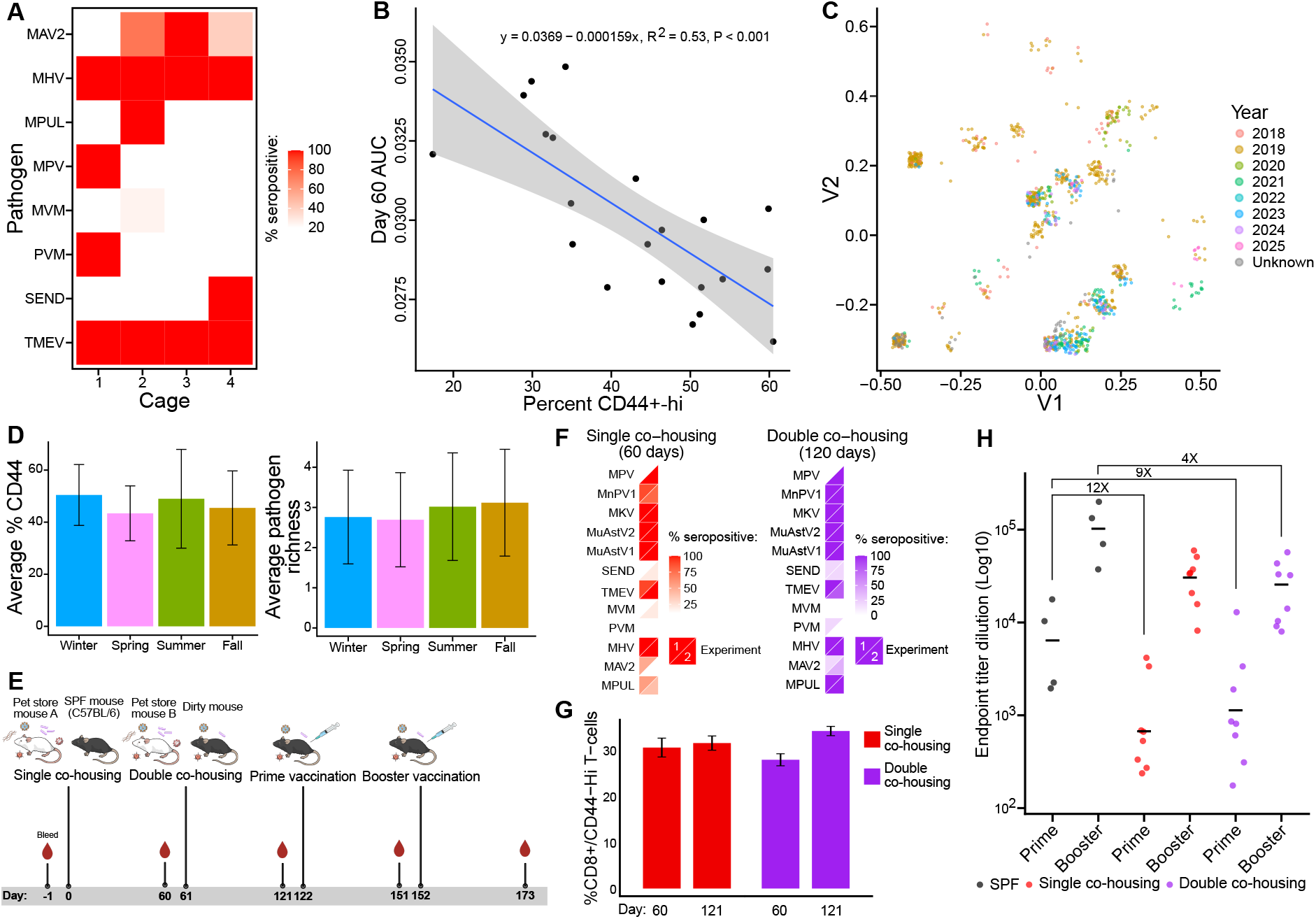
The dirty mouse model is stable over time and season. (A) Percent of mice in each cohoused cohort that were seropositive for murine pathogens. Data corresponds to mice shown in Fig. 1B-E. Only pathogens with seropositivity among the mice are depicted, additional pathogens tested in panel are listed in Materials and Methods. (B) Correlation of T cell activation status (percent of CD44+-hi expressing cells among CD8a+ T cells) with anti-S1 RBD IgG levels at 60 days post-prime (30 days post-boost). Antibody levels are represented as AUC. N=21 mice. Linear regression model equation and 95% confidence interval is shown on graph. (C) Multidimensional scaling (MDS) analysis of serology panel data from cohoused mice. Each dot represents a dirty laboratory mouse. Distance between points represents similarity of pathogen exposure history. Points are colored by the year that the experiment occurred. Data is representative of N=1014 mice. (D) Comparison of average percent CD44+-Hi among CD8a+ T cells (left) and average pathogen richness (right) from mice in panel C across seasons. Error bars represent standard deviation. (E) Experimental design used to generate the double co-housed dirty mouse model. Illustration generated with images from NIAID NIH BioArt. (F) Heatmaps of seroprevalence against a panel of murine pathogens in single co-housed (60 days) and double co-housed (120 days) mice. Squares are divided into triangles representing data from two independent experimental replicates. (G) Frequency of activated CD8a+/CD44-Hi T cells in single and double co-housed mice assessed at days 60 and 121. Bars represent mean ± SEM. (H) Anti-S1 RBD IgG log10 endpoint titers (OD_405_) 30 days post-prime and -booster (N=20) in SPF (black), single (red), and double (purple) co-housed mice. Data represent individual mice with horizontal bars indicating mean and brackets indicate fold-change relative to SPF. Antibody levels in panels B and H were determined using ELISA.

## Discussion

Despite the overwhelming success of SARS-CoV-2 mRNA vaccines at preventing hospitalization and death, efficacy of these vaccines wanes over time (15) and requires a booster vaccination to achieve optimal efficacy (13). Initial pre-clinical testing of SARS-CoV-2 mRNA vaccine candidates was performed in SPF mice and macaques raised in standard SPF conditions. Vaccination of these animals with mRNA vaccines resulted in robust neutralizing antibody titers (16, 17). Given that humans do not live in SPF conditions and regularly experience contact with pathogens, we investigated whether the humoral response in dirty mice vaccinated with SARS-CoV-2 mRNA vaccines better modelled the human response towards the vaccination. Dirty mice had lower Spike IgG serum antibody titers after the initial vaccination and experienced faster titer wane over time, and the resultant antibodies were not as effective at neutralizing WT SARS-CoV-2 or binding to omicron variants BA.1 and BA.5, as compared to SPF mice. These results are consistent with data generated in mice vaccinated with influenza virus vaccines (2) and demonstrates that co-housed mice are a valuable model for mRNA vaccines which have not been previously evaluated in dirty mice.

The mechanism underlying poorer mRNA vaccine response in dirty mice remains to be fully elucidated. Rewilding mice results in greater B cell maturation and germinal center responses (18), but it is not clear how B cell activation seems to interfere with generation of protective antibodies (2). We found that α’NP antibody cleared faster from dirty mice than SPF mice, which may in part explain the durability of the SPF vaccination response. Additional data on phenotypes of antigen-specific B cells may help elucidate whether B cells generated in response to vaccines in dirty mice are deficient in some way. The finding that T cell activation status negatively correlated with robust vaccine response suggests that immune interference is a “sliding scale”, with critical implications for clinical translation where standard vaccine dosages may eventually reach a threshold of immune maturation where they fail to protect against infection.

Pathogen exposure history is heterogeneous over time in the pet store cohousing model but yields a robust T cell activation status. This study presents a snapshot of “dirtiness” in this model over time, adding 4 years of data to our previous understanding of the robustness of the dirty mouse model (2). We found no evidence of seasonal differences in T cell activation or pathogen exposure, which may affect other methods of microbial exposure that rely on wild mice or outdoor exposure. Given the heterogeneity in pathogen exposure within and between experiments, we investigated if there were any correlations between pathogens or pathogen combinations and mRNA vaccine antibody responses. We found that pathogen number, combination, and T cell activation status were stable and comparable between 2018-2025, consistent with the stability over time of other dirty mouse platforms (19).

The basal immune signature of SPF mice closely matches that of neonatal humans (5). As individuals mature into adulthood, cumulative infections continue to shape the immune system, permanently altering memory T cell composition and expanding innate immune cell populations (1, 5). With the double co-housing model, we were able to mimic this scenario and found that pathogen recall did not affect the already established T cell activation in our model. The antibody response in the double co-housing model confirms our prior data and existing research (2, 5), where the exaggerated antibody titers typically produced in clean mice decreased as we increased the complexity of the immune system in the model.

The dirty mouse humoral response better models human SARS-CoV-2 vaccination experience, given that humans require booster doses (13, 20), experience waning serum antibody titers (20-22), and reduced variant neutralization following SARS-CoV-2 mRNA vaccination (23, 24). SPF mice are often poor predictors of human immune response, and more translationally representative dirty mice should be used for preclinical immunizations studies due to their similarity to a more developed adult human immune system.

## Acknowledgements

This work was supported by R01 AI 173043 and AI 150600 to R.A.L. F.K.S., C.A.M. and S.N.R. were supported by National Institutes of Health T32 HL007741. B.P. was supported by the University of Minnesota’s President’s Postdoctoral Fellowship Program.

This work was supported in part by grants National Institutes of Health (NIH) P51OD011132 and NIH/NIAID CEIRR under contract 75N93021C00017 to Emory University to M.S.S., Emory Executive Vice President for Health Affairs Synergy Fund award to M.S.S. and JW, the Pediatric Research Alliance Center for Childhood Infections and Vaccines and Children’s Healthcare of Atlanta to M.S.S., COVID-Catalyst-I3 Funds from the Woodruff Health Sciences Center and Emory School of Medicine to M.S.S., and Woodruff Health Sciences Center 2020 COVID-19 CURE Award to M.S.S. Funders played no role in the design and conduct of the study; collection, management, analysis, and interpretation of the data; preparation, review, or approval of the manuscript; and decision to submit the manuscript for publication.

## Supplemental methods

### Co-housing and serological analysis of mice

Age-matched female 8-to 10-week-old C57BL/6 (Jackson laboratory) mice were either housed in a specific pathogen-free (SPF) mouse facility or co-housed in a BSL-3 facility with mice acquired from a local pet store for 60 days, resulting in what is referred to as dirty mice. Thereafter, submandibular bleeding was performed on the mice to collect blood onto Opti-Spot strips (IDEXX BioAnalytics) for serological testing, whose panel tested for *Filobacterium rodentium, Encephalitozoon cuniculi* (ECUN), Ectromelia virus, Epizootic Diarrhea of Infant Mice/Mouse Rotavirus (EDIM), Lymphocytic Choriomeningitis Mammarenavirus (LCMV), *Mycoplasma pulmonis* (MPUL), Mouse Adenoviruses Type 1 (MAV1) and 2 (MAV2), Mouse Cytomegalovirus (MCMV), Mouse Hepatitis Virus (MHV), Mouse Parvovirus Type-1 (MPV), Minute Virus of Mice (MVM), Mouse Polyomavirus, Pneumonia Virus of Mice (PVM), Reovirus (REO), Theiler’s Murine Encephalomyelitis Virus (TMEV), Sendai Virus (SEND), and *Clostridium piliforme* (CPIL). In-house peptide ELISAs were developed to detect antibodies against viruses not typically included in commercial panels, specifically murine astrovirus 2 (MuAstV2), murine kobuvirus (MKV), and Minnesota rodent picornavirus 1 (MnPV1); these assays were performed as previously described (1, 2). For murine astrovirus 1 (MuAstV1), a commercial ELISA plate (XpressBio) pre-coated with recombinant capsid protein was utilized. Blood was additionally collected for PBMC (peripheral blood mononuclear cell) isolation and flow analysis for markers of immune cell identity and activation. CD8a+/CD44+-Hi cells were first gated to identify single cells, then to identify CD45+ immune cells (anti-CD45 Brilliant Violet 711 antibody, clone30F-11, BioLegend), then to identify CD8a+ cells (anti-CD8a PE/Cyanine5 antibody, clone 53-6.7, BioLegend), and finally to identify KLRG1+/CD44+-Hi cells (anti-CD44 Brilliant Violet 785 antibody, clone IM7, BioLegend; anti-KLRG1 PE/Cyanine7 antibody, clone 2F2, BioLegend). Cells were analyzed using a BD LSRFortessa™ flow cytometer (BD Biosciences), with a minimum of 100,000 live events acquired per sample. For the double co-housing model, a subset of the dirty mice from the initial 60-day exposure was subjected to a second 60-day co-housing period (double Co-Ho) prior to immunization. The Institutional Animal Care and Use Committee (IACUC) of the University of Minnesota approved all animal experimentation performed in this work.

### Vaccination/serum collection

Mice were vaccinated intramuscularly in the rear leg with either 0.25 µg Pfizer-BioNTech COVID-19 vaccine (PAA165969, Lot EN6025) or with 0.5 µg of mRNA encoding the SARS-CoV-2 receptor-binding domain (RBD) derived from the WA1 variant (2020 SARS-CoV-2 A.1 strain mRNA vaccine), formulated in lipid nanoparticles, and delivered in PBS for a total volume of 50 µl. The vaccine design was in accordance with World Health Organization (WHO) recommendations for variant-based COVID-19 immunogens. Vaccination occurred 61-113 days post-co-housing in 24-week-old female mice. Mice were primed (day 0) and provided an identical booster vaccination after 31 days. For the double co-housing, mice were primed on day 120 post-co-housing, corresponding to 60 days after completion of the first co-housing protocol, and received a homologous booster immunization 31 days later. Submandibular bleedings were performed on the days immediately prior to vaccination (d0), 30 days post-prime prior to the booster dose (d30), 30 days post-boost (d61-62), and monthly subsequent to the booster vaccination (d91, d122, and d151) until the end of the experiment. Serum was isolated from collected blood following one hour of coagulation at room temperature, followed by 15 minutes’ centrifugation at 3,000 x G, 4ºC. Serum was stored at -80ºC until experimental use.

### Vaccine-specific antibody detection

Vaccine-specific antibodies were detected with either ELISA or the 10-plex Meso Scale Discovery-Electrochemiluminescence immunoassay (MSD-ECLIA), as stated in figure legends. For ELISA, 96-well plates (Corning Inc.) were coated with 3 µg/mL of recombinant SARS-CoV-2 S1 RBD protein (3) diluted in phosphate-buffered saline (PBS; Bio-Rad Laboratories, Inc.) overnight at 4ºC. The following day, plates were washed three times with PBS containing 0.5% Tween-20 (PBS-T; Hoefer, Inc.), then blocked with 1% bovine serum albumin (BSA; MP Biomedicals) in PBS for 1 hour at room temperature to prevent nonspecific binding. After blocking, plates were washed again three times with PBS-T. Mouse serum was diluted 1:50 in dilution buffer (PBS containing 0.05% Tween-20 and 0.2% BSA), then serially diluted 1:4 in dilution buffer, added to the plates, incubated for 1 hour at 37ºC, and subsequently washed three times with PBS-T. Bound serum antibodies were detected by incubating the plate for 1 hour at 37ºC with horseradish peroxidase (HRP)-conjugated anti-mouse IgG antibodies (Southern Biotech) diluted 1:500 in blocking buffer, and the plate was subsequently washed four times in PBS-T. Plates were then incubated with ABTS peroxidase substrate (SeraCare) for 45 minutes at room temperature in the dark, and reaction was stopped with 2% sodium dodecyl sulfate (SDS; Fisher Scientific) in PBS. For visualization, optical density at 405 nm (OD405) was measured using a Synergy H1 microplate reader (BioTek). OD405 values were plotted against the dilution factor, and the area under the curve (AUC) was calculated using GraphPad Prism 10.

For MSD-ECLIA (Rockville, MD), SARS-Cov2 spike IgG was quantified using plates coated with WA.1, BA.1, or BA.5 spike proteins (4). Briefly, plates were blocked with 5% BSA for 60 minutes prior to the addition of samples diluted at 1:5000 and allowed to incubate with shaking for two hours. Then, plates were washed and labeled with Sulfo-tagged anti-IgG antibody. Plates were washed to remove unbound detection antibodies and binding was detected via electrochemiluminescence on an MSD Sector instrument after addition of ECL substrate. Raw data were analyzed using Discovery Workbench and GraphPad Prism software v10.1.2. The antibody concentrations were calculated relative to antibody standard controls and expressed in arbitrary units (AU) per mL for IgG.

### Cells and Viruses

VeroE6-TMPRSS2 cells were generated and cultured as previously described (5). All SARS-CoV-2 viruses were plaque purified and propagated once in VeroE6-TMPRSS2 cells to generate working stocks. Viruses were deep sequenced and confirmed as previously described (6).

### Focus Reduction Neutralization Test (FRNT)

FRNT assays were performed as previously described (5-7). Briefly, serum samples were 3-fold diluted in 8 serial dilutions using DMEM with an initial dilution of 1:10. Serially diluted samples were mixed with an equal volume of the desired SARS-CoV-2 variant (100-200 foci per well). The virus-serum mixtures were incubated at 37°C for 1 hour in a round bottom 96-well culture plate. After 1 hour incubation, the virus-serum mixture was added to VeroE6-TMPRSS2 cells and incubated at 37°C for an additional hour. Post-incubation, the mixture was removed from cells and 100μl of prewarmed 0.85% methylcellulose overlay was added to each well. Plates were incubated at 37°C for 18 to 40 hours (depending on the SARS-CoV-2 strain). After the appropriate incubation time, the methylcellulose overlay was removed, and cells were washed with PBS and fixed with 2% paraformaldehyde for 30 minutes. Following fixation, cells were washed twice with PBS, and permeabilized using permeabilization buffer for at least 20 minutes. After permeabilization, cells were incubated with an anti-SARS-CoV-2 spike primary antibody directly conjugated to Alexa Fluor-647 (CR3022-AF647) overnight at 4°C. Cells were then washed twice with 1X PBS and imaged on an ELISPOT reader (CTL Analyzer).

### Seasonality analysis

Serology testing results from this study were combined with historical serology panel testing data from dirty mice. Historical data included C57BL/6 female mice that were cohoused with pet store mice, Balb/c female mice that were cohoused with pet store mice, and C57BL/6 male mice that were exposed to pet store mice fomites (male mice cannot be cohoused with pet store mice due to ethical concerns). Historical data was collected from 2018-2024 and includes previously published serology data (8). In total, 1014 data points were included in the analysis. Serology panel data were combined and limited to the pathogens shared in all panel assays and included CPIL, ECUN, EDIM, TMEV, LCMV, MAV, MCMV, MHV, MPUL, MPV, MVM, POLY, PVM, REO, and SEND. The IDEXX panel used for this work included separate tests for MAV1 and MAV2, and these tests were collapsed to a single MAV result (if mice were positive for either MAV1 or MAV2, they were considered MAV-positive). Historical panels tested for several serotypes of MPV in separate, and these data were similarly collapsed to a single MPV result to combine with the panel data used in this study (if mice were positive for any of the MPV targets, they were considered MPV-positive). Serology panel data (binary data of either seropositive or seronegative) were used for multidimensional scaling analysis using the stats package in R (9). The percentage of CD8a+/CD44+-Hi T cells were compared across the 1014 mice to identify differences in seasonality of T cell activation status. The number of seropositive pathogens per mouse was also compared across seasons.

### Statistical analyses

To compare rates of vaccine-specific antibody decline (Fig. 1C), a linear regression was fit to SARS-CoV-2 WT spike IgG levels (AU/ml) for the dirty and SPF mice for data collected at 62-, 91-, 122-, and 151-days post-priming. The slopes were compared using Welch’s T test. The percentage of CD8a+/CD44+-Hi T cells was correlated with antibody levels (measured by AUC) on day 60 (30 days-post boost) using Pearson correlation. Antibody neutralization was quantified by counting the number of foci for each sample using the Viridot program (10). The neutralization titers were calculated as follows: 1 – (ratio of the mean number of foci in the presence of sera and foci at the highest dilution of the respective sera sample). Each sample was tested in duplicate. The FRNT-50 titers were interpolated using a 4-parameter nonlinear regression in GraphPad Prism 10.1.2. Samples that do not neutralize at the limit of detection at 50% are plotted at 20 and used for geometric mean and fold-change calculations.

### Antibody clearance

Mice were tail vein-injected either with 200 µg influenza A nucleoprotein-targeting (α’NP) antibody (clone H16-L10-4R5 (HB-65); BioXCell) brought to 100 µl total with InVivo Pure Ph 7.0 dilution buffer (BioXCell), or with 100 µl dilution buffer alone for the mock condition. In total, three mock SPF mice, four α’NP-injected SPF mice, and four α’NP-injected dirty mice were used. Submandibular bleedings of mice were performed one week prior to injection (d-7), the day following injection (d1), and 2, 4, 6, 8, 12, and 17 weeks after injection (d14, d29, d43, d59, d90, d120). To assess serum α’NP antibody levels, ELISA assays were performed as previously described on plates coated with 1 µg/ml His-tagged Influenza A H1N1 (A/Puerto Rico/8/34/Mount Sinai) Nucleoprotein (I116M; Sino Biological, Inc.) diluted in PBS. Serum was diluted 1:100 in dilution buffer prior to serial 1:4 dilutions, and Goat Anti-Mouse IgG2a Human ads-HRP secondary antibody (SouthernBiotech) was used as the secondary antibody for detection. AUC and standard deviation per timepoint were calculated and plotted using GraphPad Prism 10.

### Figure preparation

The following NIH BioArt (11) images were used or modified to generate Figure 1A and Figure 2E: BIOART-000042, BIOART-000279, BIOART-000464, BIOART-000496, BIOART-000505, BIOART-000646, and BIOART-000652.

### Data and code availability

Code for MDS analysis of pathogen exposure is located on github at github.com/langloislab/praena-shepherd-mcdonald-2026.

